## Supplemental Data 1 for "Amis *Pacilo* and Yami *Cipoho* are not the same as the Pacific breadfruit starch crop—Target enrichment phylogenomics of a long-misidentified *Artocarpus* species sheds light on the northward Austronesian migration from the Philippines to Taiwan"

### S1 Appendix

#### List of literature of the Taiwanese 'breadfruit' and the misused scientific names.

#### *Artocarpus altilis* (Parkinson) Fosberg.


1. Liu T-S. Illustrations of Native and Introduced Ligneous Plants of Taiwan, Vol. 2. Taipei: College of Agriculture, National Taiwan University; 1962.
2. Liu T-S, Liao J-C. Moraceae. In: Li H-L, Liu T-S, Huang T-C, Koyama T, DeVol CE, editors. Flora of Taiwan, Vol 2. Taipei, Taiwan: Epoch Publishing Co.; 1976. p. 117–61.
3. Ying S-S. Coloured Illustrated Flora of Taiwan, Vol 3. Taipei: Published by the author; 1988.
4. Chen C-C, Huang Y-L, Ou J-C, Lin C-F, Pan T-M. Three new prenylflavones from *Artocarpus altilis*. J Nat Prod. 1993; 56(9):1594–7.
5. Hu B-Y, Hsiao W-W, Fu C-H. First report of zonate leaf spot of *Artocarpus altilis* caused by *Cristulariella moricola* in Taiwan. Plant Dis. 2002; 86(10):1179.
6. Wang H-H, Chang L-W, Kao R-C. Forest management of aborigines Tao in Lanyu (Botel Tabago) and its effects to forest structure and species composition. J Natl Park. 2003; 13(1):75–94.
7. Lan W-C, Tzeng C-W, Lin C-C, Yen F-L, Ko H-H. Prenylated flavonoids from *Artocarpus altilis*: Antioxidant activities and inhibitory effects on melanin production.


#### *Artocarpus communis* J.R.Forst. & G.Forst. [=*A. altilis* (Parkinson) Fosberg]


8. Sasaki S. List of Plants of Formosa. Taihoku: Natural History Society of Formosa; 1928.
9. Kudô Y. Iconography of Tropical Plants in Taiwan, Vol. 1. Tokyo: Meibundo & Co.; 1934.
10. Yang C-F. Manual of Fruit Trees in Taiwan. Chiayi: Chiayi Experiment Station, Taiwan Agricultural Research Institute; 1951.
11. Lin CN, Shieh WL. Prenylflavonoids and a pyranodihydrobenzoxanthone from *Artocarpus communis*. Phytochemistry. 1991; 30(5):1669–71.
12. Shieh W-L, Lin C-N. A quinonoid pyranobenzoxanthone and pyranodihydrobenzoxanthone from *Artocarpus communis*. Phytochemistry. 1992; 31(1):364–7.
13. Lin CN, Shieh WL, Jong TT. A pyranodihydrobenzoxanthone epoxide from *Artocarpus communis*. Phytochemistry. 1992; 31(7):2563–4.
14. Lin CN, Shieh WL. Pyranoflavonoids from *Artocarpus communis*. Phytochemistry. 1992; 31(8):2922–4.
15. Lin C-N, Shieh W-L, Ko F-N, Teng C-M. Antiplatelet activity of some prenylflavonoids. Biochem Pharmacol. 1993; 45(2):509–12.
16. Liou S-S, Shieh W-L, Cheng T-H, Won S-J, Lin C-N. γ-pyrone compounds as potential anti-cancer drug. J Pharm Pharmacol. 1993; 45(9):791–4.
17. Lin C-C. Moraceae. In: Yang Y-P, Liu H-Y, Lu S-Y, editors. Manual of Taiwan Vascular Plants, Vol 2. Taipei: The Council of Agriculture, The Executive Yuan; 1999. p. 46–57.
18. Cheng H-W, Lu S-Y. Botel Tabaco, Yami & Plants. Taipei: Lamper Enterprises Co., Ltd.; 2000.
19. Chan S-C, Ko H-H, Lin C-N. New prenylflavonoids from *Artocarpus communis*. J Nat Prod. 2003; 66(3):427–30.
20. Wei B-L, Weng J-R, Chiu P-H, Hung C-F, Wang J-P, Lin C-N. Antiinflammatory flavonoids from *Artocarpus heterophyllus* and *Artocarpus communis*. J Agr Food Chem. 2005; 53(10):3867–71.
21. Weng J-R, Chan S-C, Lu Y-H, Lin H-C, Ko H-H, Lin C-N. Antiplatelet prenylflavonoids from *Artocarpus communis*. Phytochemistry. 2006; 67(8):824–9.
22. Fang S-C, Hsu C-L, Yu Y-S, Yen G-C. Cytotoxic effects of new geranyl chalcone derivatives isolated from the leaves of *Artocarpus communis* in SW 872 human liposarcoma cells. J Agr Food Chem. 2008; 56(19):8859–68.
23. Lin K-W, Liu C-H, Tu H-Y, Ko H-H, Wei B-L. Antioxidant prenylflavonoids from *Artocarpus communis* and

*Artocarpus elasticus*. Food Chem. 2009; 115(2):558–62.
24. Ko H-H, Lan W-C, Zhan W-Y. Inhibitory effects of prenyl flavonoids isolated from *Artocarpus communis* on mushroom tyrosinase and melanin production in cultured melanoma cells. Planta Med. 2010; 76(12):1248.
25. Hsu C-L, Shyu M-H, Lin J-A, Yen G-C, Fang S-C. Cytotoxic effects of geranyl flavonoid derivatives from the fruit of *Artocarpus communis* in SK-Hep-1 human hepatocellular carcinoma cells. Food Chem. 2011; 127(1):127–34.
26. Lin J-A, Fang S-C, Wu C-H, Huang S-M, Yen G-C. Anti-inflammatory effect of the 5,7,4 '-trihydroxy-6-geranylflavanone Isolated from the fruit of *Artocarpus communis* in S100B-Induced human monocytes. J Agr Food Chem. 2011; 59(1):105–11.
27. Hsu C-L, Chang F-R, Tseng P-Y, Chen Y-F, El-Shazly M, Du Y-C, et al. Geranyl flavonoid derivatives from the fresh leaves of *Artocarpus communis* and their anti-inflammatory activity. Planta Med. 2012; 78(10):995–1001.
28. Lin J-A, Wu C-H, Fang S-C, Yen G-C. Combining the observation of cell morphology with the evaluation of key inflammatory mediators to assess the anti-inflammatory effects of geranyl flavonoid derivatives in breadfruit. Food Chem. 2012; 132(4):2118–25.
29. Ko H-H, Tsai Y-T, Yen M-H, Lin C-C, Liang C-J, Yang T-H, et al. Norartocarpetin from a folk medicine *Artocarpus communis* plays a melanogenesis inhibitor without cytotoxicity in B16F10 cell and skin irritation in mice. BMC Complem Altern Med. 2013; 13:e348.
30. Lee CW, Ko HH, Chai CY, Chen WT, Lin CC, Yen FL. Effect of *Artocarpus communis* extract on UVB irradiation-induced oxidative stress and inflammation in hairless mice. Int J Mol Sci. 2013; 14(2):3860–73.
31. Lee C-W, Ko H-H, Lin C-C, Chai C-Y, Chen W-T, Yen F-L. Artocarpin attenuates ultraviolet B-induced skin damage in hairless mice by antioxidant and anti-inflammatory effect. Food Chem Toxicol. 2013; 60:123–9.
32. Fu Y-T, Lee C-W, Ko H-H, Yen F-L. Extracts of *Artocarpus communis* decrease α-melanocyte stimulating hormone-induced melanogenesis through activation of ERK and JNK signaling pathways. Sci World J. 2014; 2014:e724314.
33. Lin J-A, Chen H-C, Yen G-C. The preventive role of breadfruit against inflammation-associated epithelial carcinogenesis in mice. Mol Nutr Food Res. 2014; 58(1):206–10.
34. Tzeng C-W, Yen F-L, Lin L-T, Lee C-W, Yen M-H, Tzeng W-S, et al. Antihepatoma activity of *Artocarpus communis* is higher in fractions with high artocarpin content. Sci World J. 2014; 2014:e978525.
35. Hu SC-S, Lin C-L, Cheng H-M, Chen G-S, Lee C-W, Yen F-L. Artocarpin induces apoptosis in human cutaneous squamous cell carcinoma HSC-1 cells and its cytotoxic activity is dependent on protein-nutrient concentration. Evid-Based Compl Alt Med. 2015; 2015:e236159.
36. Lin K-W, Wang B-W, Wu C-M, Yen M-H, Wei B-L, Hung C-F, et al. Antioxidant prenylated phenols of *Artocarpus* plants attenuate ultraviolet radiation-induced damage on human keratinocytes and fibroblasts. Phytochem Lett. 2015; 14:190–7.
37. Tzeng C-W, Tzeng W-S, Lin L-T, Lee C-W, Yen M-H, Yen F-L, et al. *Artocarpus communis* induces autophagic instead of apoptotic cell death in human hepatocellular carcinoma cells. Am J Chinese Med. 2015; 43(3):559-79.
38. Tzeng C-W, Tzeng W-S, Lin L-T, Lee C-W, Yen F-L, Lin C-C. Enhanced autophagic activity of artocarpin in human hepatocellular carcinoma cells through improving its solubility by a nanoparticle system. Phytomedicine. 2016; 23(5):528-40.
39. Chung S-W. Illustrated Flora of Taiwan, Vol. 4. Taipei: Owl Publishing House Co., Ltd.; 2017.
40. Lin J-A, Wu C-H, Yen G-C. Breadfruit flavonoid derivatives attenuate advanced glycation end products (AGEs)-enhanced colon malignancy in HCT116 cancer cells. J Funct Foods. 2017; 31:248–54.
41. Tsai M-H, Liu J-F, Chiang Y-C, Hu SC-S, Hsu L-F, Lin Y-C, et al. Artocarpin, an isoprenyl flavonoid, induces p53-dependent or independent apoptosis via ROS-mediated MAPKs and Akt activation in non-small cell lung cancer cells. Oncotarget. 2017; 8(17):28342–58.
42. Yeh C-J, Chen C-C, Leu Y-L, Lin M-W, Chiu M-M, Wang S-H. The effects of artocarpin on wound healing:

in vitro and in vivo studies. Sci Rep. 2017; 7:e15599.

43. Lee CW, Hsu LF, Lee MH, Lee IT, Liu JF, Chiang YC, et al. Extracts of *Artocarpus communis* induce mitochondria-associated apoptosis via pro-oxidative activity in human glioblastoma cells. Front Pharmacol. 2018; 9:e411.
44. Yang C-Y, Huang P-H, Tseng C-H, Yen F-L. Topical *Artocarpus communis* nanoparticles improved the water solubility and skin permeation of raw *A. communis* extract, improving its photoprotective effect. Pharmaceutics. 2021; 13(9):e1372.

### *Artocarpus incisus* (Thunb.) L.f. [=*A. altilis* (Parkinson) Fosberg]

45. Mackay GL. From Far Formosa: The Islands, its People and Mission. MacDonald JA, editor. New York, Chicago and Toronto: Fleming H. Revell Company; 1896.
46. Kawakami T. A List of Plants of Formosa. Taihoku: Bureau of Productive Industry, Government of Formosa; 1910.
47. Hayata B. Materials for a Flora of Formosa. J Coll Sci Imp Univ Tokyo. 1911; 30(1):1–471.
48. Sasaki S. Important plants of Botel Tobago. J Nat Hist Soc Taiwan. 1913; 3(9):35–43.
49. Tashiro Y. A guide to street trees and planting trees in urban Taiwan. Taihoku: Bureau of Forest Management, Government of Formosa; 1920.
50. Chang C-E. An enumeration of the woody plants of Botel Tobago. J Phytogeogr Taxon. 1981; 29(1):1–21.
51. Yamazaki T. Taxonomic review of the Moraceae from Japan, Korea, Taiwan and adjacent areas (1). J Phytogeogr Taxon. 1982; 30(2):61-73.
52. Liao J-C. A taxonomic revision of the family Moraceae in Taiwan (1) Genera *Artocarpus*, *Broussonetia* and *Fatoua*. Quart J Exp For Natl Taiwan Univ. 1989; 3(1):145–51.
53. Liu Y-C, Lu F-Y, Ou C-H. Trees of Taiwan, Revised edition. Taichung: College of Agriculture, National Chung-Shing University; 1994.
54. Liao J-C. Moraceae. In: Editorial Committee of the Flora of Taiwan, editor. Flora of Taiwan, Vol 2, 2nd edn. Taipei, Taiwan: Department of Botany, National Taiwan University; 1996. p. 136–95.
55. Wang H-H, Chang L-W, Kao R-C. Forest management of aborigines Tao in Lanyu (Botel Tobago) and its effects to forest structure and species composition. J Natl Park. 2003; 13(1):75–94.
56. Yang C-K, Jung M-C. An ethnobotanical Memory of Taiwan. Taichung: Morning Star Publishing Inc.; 2012.
57. Editorial Committee of the Red List of Taiwan Plants. The Red List of Vascular Plants of Taiwan, 2017: Endemic Species Research Institute, Forestry Bureau, Council of Agriculture, Executive Yuan and Taiwan Society of Plant Systematics; 2017.

### *Artocarpus integrifolia* L. (=*A. heterophyllus* Lam.)

58. Tashiro Y. A guide to planting trees in urban Taiwan. Taihoku: Bureau of Productive Industry, Government of Formosa; 1900.
59. Matsumura J, Hayata B. Enumeratio plantarum in insula Formosa sponte crescentium hucusque rite cognitarum adjectis descriptionibus et figuris specierum pro regione novarum. J Coll Sci Imp Univ Tokyo. 1906; 22:1–702.

### *Artocarpus lanceolatus* Trécul (=*A. lamellosus* Blanco)

60. Li H-L. Woody Flora of Taiwan. Narberth, Pennsylvania: Livingston Publishing Company; 1963.
