## Supplementary material for "Amis *Pacilo* and Yami *Cipoho* are not the same as the Pacific breadfruit starch crop—Target enrichment phylogenomics of a long-misidentified *Artocarpus* species sheds light on the northward Austronesian migration from the Philippines to Taiwan": S1 Table. Summary statistics of assembly quality of the 64 samples used in this study.

| SRAAccession | CollectionNo | NumReads | ReadsMapped | PctOnTarget | GenesMapped | GenesWithContigs | GenesWithSeqs | GenesAt25pct | GenesAt50pct | GenesAt75pct | Genesat150pct | ParalogWarnings |
|---|---|---|---|---|---|---|---|---|---|---|---|---|
| SRR19994412 | CCR089 | 3454253 | 2509285 | 0.726 | 525 | 524 | 523 | 523 | 520 | 498 | 0 | 59 |
| SRR19994411 | CCR114 | 2465844 | 1588078 | 0.644 | 526 | 524 | 524 | 524 | 521 | 499 | 1 | 52 |
| SRR19994400 | CCR120 | 3372355 | 2296884 | 0.681 | 527 | 524 | 523 | 523 | 520 | 502 | 2 | 59 |
| SRR19994389 | CCR123 | 1598087 | 989733 | 0.619 | 526 | 524 | 524 | 523 | 519 | 488 | 3 | 40 |
| SRR19994382 | CCR127 | 4896763 | 3472019 | 0.709 | 526 | 524 | 523 | 523 | 521 | 504 | 1 | 67 |
| SRR19994381 | CCR129 | 4819831 | 3500315 | 0.726 | 528 | 524 | 523 | 523 | 522 | 506 | 1 | 70 |
| SRR19994380 | CCR131 | 5478720 | 3837157 | 0.7 | 527 | 524 | 523 | 522 | 522 | 504 | 0 | 62 |
| SRR19994379 | CCR134 | 4227752 | 2979424 | 0.705 | 526 | 524 | 523 | 523 | 521 | 500 | 0 | 66 |
| SRR19994378 | CCR146 | 4403105 | 3188458 | 0.724 | 526 | 524 | 523 | 523 | 522 | 505 | 1 | 66 |
| SRR19994377 | CCR155 | 5086908 | 3501802 | 0.688 | 528 | 524 | 523 | 523 | 522 | 504 | 0 | 68 |
| SRR19994410 | CCR161 | 5578338 | 3961632 | 0.71 | 526 | 525 | 524 | 523 | 522 | 506 | 2 | 75 |
| SRR19994409 | CCR168 | 4612233 | 3253168 | 0.705 | 525 | 524 | 524 | 524 | 523 | 506 | 1 | 69 |
| SRR19994408 | CCR172 | 5387868 | 3809024 | 0.707 | 526 | 524 | 524 | 524 | 522 | 505 | 0 | 67 |
| SRR19994407 | CCR174 | 7074427 | 5066260 | 0.716 | 526 | 525 | 524 | 524 | 523 | 506 | 0 | 77 |
| SRR19994406 | CCR176 | 4923132 | 3370044 | 0.685 | 525 | 524 | 523 | 523 | 522 | 505 | 0 | 75 |
| SRR19994405 | CCR178 | 5594222 | 3897376 | 0.697 | 525 | 524 | 523 | 523 | 522 | 504 | 0 | 71 |
| SRR19994404 | CCR179 | 4977478 | 3468262 | 0.697 | 526 | 524 | 523 | 523 | 522 | 504 | 0 | 70 |
| SRR19994403 | CCR182 | 5938451 | 4222404 | 0.711 | 525 | 524 | 523 | 523 | 522 | 506 | 0 | 71 |
| SRR19994402 | CCR184 | 4751564 | 3262344 | 0.687 | 527 | 525 | 524 | 524 | 523 | 505 | 0 | 68 |
| SRR19994401 | CCR195 | 6224964 | 4276762 | 0.687 | 527 | 524 | 523 | 523 | 522 | 506 | 0 | 70 |
| SRR19994399 | CCR198 | 7061771 | 4933917 | 0.699 | 528 | 524 | 524 | 524 | 523 | 506 | 0 | 77 |
| SRR19994398 | CCR200 | 5875516 | 4031108 | 0.686 | 530 | 530 | 530 | 530 | 529 | 513 | 1 | 56 |
| SRR19994397 | CCR202 | 6288263 | 4473014 | 0.711 | 526 | 524 | 524 | 524 | 523 | 506 | 0 | 74 |
| SRR19994383 | MYT20 | 4906883 | 3323339 | 0.677 | 529 | 521 | 521 | 520 | 519 | 501 | 2 | 101 |
| SRR19994385 | CHW72 | 5351261 | 3821967 | 0.714 | 526 | 525 | 524 | 524 | 523 | 505 | 1 | 69 |
| SRR19994384 | CHW75 | 5274716 | 3728560 | 0.707 | 527 | 525 | 524 | 524 | 523 | 506 | 1 | 68 |
| SRR19994396 | CCR203 | 5144804 | 3607582 | 0.701 | 527 | 524 | 523 | 523 | 522 | 506 | 1 | 69 |
| SRR19994395 | CCR204 | 6849580 | 5075154 | 0.741 | 524 | 515 | 512 | 509 | 502 | 478 | 1 | 98 |
| SRR19994394 | CCR205 | 5129715 | 3579730 | 0.698 | 527 | 524 | 524 | 524 | 522 | 506 | 0 | 71 |
| SRR19994393 | CCR207 | 4499613 | 3249875 | 0.722 | 525 | 518 | 516 | 513 | 509 | 493 | 0 | 102 |
| SRR19994392 | CCR208 | 5101754 | 3637717 | 0.713 | 528 | 526 | 525 | 525 | 523 | 504 | 1 | 67 |
| SRR19994391 | CCR209 | 7324117 | 5264746 | 0.719 | 528 | 525 | 524 | 524 | 523 | 505 | 0 | 75 |
| SRR19994390 | CCR217 | 7080167 | 5058343 | 0.714 | 530 | 527 | 527 | 526 | 524 | 508 | 0 | 69 |
| SRR19994388 | CCR218 | 6528776 | 4652004 | 0.713 | 528 | 524 | 523 | 523 | 522 | 506 | 0 | 74 |
| SRR19994387 | CCR219 | 6874864 | 4989097 | 0.726 | 527 | 524 | 524 | 524 | 522 | 506 | 0 | 70 |
| SRR19994386 | CCR220 | 7911324 | 5651708 | 0.714 | 527 | 526 | 526 | 526 | 524 | 505 | 1 | 75 |
| SRR12282879 | K7 | 560727 | 315276 | 0.562 | 530 | 507 | 507 | 507 | 507 | 492 | 1 | 36 |
| SRR12282882 | PPI2741 | 3841988 | 2379077 | 0.619 | 529 | 401 | 382 | 352 | 243 | 142 | 0 | 1 |
| SRR12282885 | Elmer 16247 | 4326990 | 1504127 | 0.348 | 521 | 449 | 439 | 408 | 291 | 140 | 0 | 0 |
| SRR12282890 | Escritor | 4411842 | 679277 | 0.154 | 528 | 458 | 417 | 372 | 253 | 126 | 0 | 0 |
| SRR12282895 | PPI3911 | 406084 | 52930 | 0.13 | 505 | 212 | 190 | 100 | 18 | 1 | 0 | 0 |
| SRR12282900 | S26230 | 2232808 | 985185 | 0.441 | 523 | 463 | 460 | 425 | 348 | 222 | 0 | 0 |
| SRR12282901 | PPI2376 | 4879436 | 2013486 | 0.413 | 528 | 511 | 510 | 506 | 471 | 396 | 2 | 12 |
| SRR12282902 | Ramos BS 42018 | 7399156 | 3427209 | 0.463 | 530 | 525 | 525 | 524 | 521 | 486 | 2 | 25 |
| SRR12282984 | Elmer13135 | 393690 | 76813 | 0.195 | 525 | 343 | 330 | 289 | 188 | 90 | 0 | 0 |
| SRR12282985 | Elmer12468 | 265445 | 89621 | 0.338 | 524 | 289 | 277 | 226 | 120 | 50 | 0 | 0 |
| SRR12282995 | Ramos BS 34736 | 284979 | 212769 | 0.747 | 525 | 452 | 447 | 423 | 339 | 208 | 1 | 1 |
| SRR12283033 | Yang 15648 | 1090597 | 800220 | 0.734 | 525 | 505 | 501 | 499 | 481 | 453 | 1 | 67 |
| SRR12283036 | Yang 13056 | 643170 | 451759 | 0.702 | 528 | 515 | 515 | 515 | 511 | 476 | 0 | 45 |
| SRR12283044 | RM 45 | 253522 | 195477 | 0.771 | 506 | 454 | 452 | 445 | 411 | 353 | 2 | 17 |
| SRR12283047 | E. Gardner & al. 294 | 408393 | 256627 | 0.628 | 525 | 485 | 483 | 481 | 477 | 452 | 1 | 67 |
| SRR12283058 | N. Zerega & al. 203 | 1982713 | 1313959 | 0.663 | 529 | 525 | 524 | 524 | 522 | 498 | 2 | 58 |
| SRR12283061 | N. Zerega & al. 61 | 882700 | 557655 | 0.632 | 516 | 494 | 488 | 485 | 474 | 451 | 1 | 68 |
| SRR12283088 | E. Gardner & al. 411 | 58872 | 37544 | 0.638 | 490 | 251 | 241 | 237 | 214 | 155 | 0 | 5 |
| SRR12283097 | E. Gardner & al. 429 | 592471 | 355347 | 0.6 | 529 | 507 | 507 | 507 | 505 | 490 | 1 | 44 |
| SRR12283098 | MV2 | 1455752 | 724204 | 0.497 | 530 | 519 | 517 | 517 | 517 | 503 | 1 | 29 |
| SRR12283102 | V8 | 998464 | 605442 | 0.606 | 530 | 525 | 525 | 525 | 525 | 509 | 1 | 46 |
| SRR12283115 | N. Zerega & al. 618 | 557429 | 327804 | 0.588 | 524 | 496 | 495 | 494 | 492 | 478 | 1 | 79 |
| SRR12283118 | M10 | 614003 | 342628 | 0.558 | 529 | 512 | 511 | 511 | 510 | 494 | 2 | 28 |
| SRR12283119 | McB1 | 1110941 | 654084 | 0.589 | 529 | 527 | 527 | 527 | 527 | 515 | 1 | 47 |
| SRR15903816 | PPI1915 | 1057108 | 642679 | 0.608 | 530 | 390 | 313 | 268 | 128 | 41 | 0 | 0 |
| SRR3907129 | N. Zerega & al. 609 | 1018527 | 574894 | 0.564 | 512 | 494 | 490 | 490 | 484 | 466 | 1 | 84 |
| SRR3907333 | NZ866 | 460695 | 243475 | 0.528 | 523 | 507 | 506 | 505 | 500 | 486 | 3 | 82 |
| SRR3907497 | E. Gardner & al. 98 | 1192814 | 678220 | 0.569 | 524 | 506 | 506 | 504 | 501 | 485 | 1 | 117 |
